# Active Notch Signaling is Required for Arm Regeneration in a Brittle Star

**DOI:** 10.1101/2019.12.13.875401

**Authors:** Vladimir Mashanov, Jennifer Akiona, Maleana Khoury, Jacob Ferrier, Robert Reid, Denis Jacob Machado, Olga Zueva, Daniel Janies

**Affiliations:** Department of Biology, University of North Florida, Jacksonville, FL, USA; Wake Forest Institute for Regenerative Medicine, Winston Salem, NC, USA; University of North Carolina at Charlotte, Charlotte, NC, USA

**Keywords:** regeneration, Notch pathway, Echinoderms, gene expression, RNA-Seq, signaling pathway

## Abstract

Cell signaling pathways play vital roles in coordinating cellular events in development. For example, the Notch signaling pathway is highly conserved across all multicellular animals and is known to coordinate a multitude of diverse cellular events, including proliferation, differentiation, fate specification, and cell death. Specific functions of the pathway are, however, highly context-dependent and are not well characterized in post-traumatic regeneration. Here, we use a small-molecule inhibitor of the pathway (DAPT) to demonstrate that Notch signaling is required for proper arm regeneration in the brittle star *Ophioderma brevispina*, a highly regenerative member of the phylum Echinodermata. We also employ a transcriptome-wide gene expression analysis to characterize the downstream genes controlled by the Notch pathway in the brittle star regeneration. We demonstrate that arm regeneration involves an extensive crosstalk between the Notch pathway and other cell signaling pathways. In the regrowing arm, Notch regulates the composition of the extracellular matrix, cell migration, proliferation, and apoptosis, as well as components of the innate immune response. We also show for the first time that Notch signaling regulates the activity of several transposable elements. Our data also suggests that one of the possible mechanisms through which Notch sustains its activity in the regenerating tissues is via suppression of Neuralized1.

## Introduction

In metazoans, a surprisingly small number of signaling pathways are required to control animal development (1). Knowledge of how these pathways work and interact in different contexts is key to gaining a mechanistic understanding of processes, including developmental growth and post-traumatic regeneration. Post-traumatic regeneration often requires dynamic changes in the balance between undifferentiated progenitors and specialized differentiated cells. In response to trauma, the cells of the damaged adult tissue have to be activated and instructed to re-enter the cell cycle, engage in migratory behavior with coordinated spatial rearrangements, and eventually, differentiate into specialized cell types of a new body part. As regeneration progresses, most of the newly generated cells have to stop dividing and migrating to differentiate into specialized cell types or enter a quiescent state. A proportion of the cells are spared from terminal differentiation to be able to serve as a source of new cells for normal cell turnover or future regeneration events. Therefore, regeneration requires complex changes in cell dynamics, which must be tightly coordinated in space and time by genetically encoded signaling pathways.

The genes encoding for the Notch signaling pathway are highly conserved in the animal kingdom (1–4). This pathway regulates various key cellular events including differentiation, fate specification, proliferation, death, and patterning into tissues (2–8). The functional role of Notch signaling has been extensively characterized in both vertebrate and invertebrate embryos, as well as in adult tissues that undergo dynamic cell turnover. Functions of the Notch signaling are very diverse, context-dependent, and often even opposing. Depending on the particular organisms and the tissue, Notch can either promote cell differentiation or facilitate stem cell maintenance, suppress or facilitate cancer progression, and induce synthesis or degradation of the extracellular matrix components (5, 8).

The Notch signaling pathway works through a juxtacrine mechanism. Both Notch ligands (on the target cell) and receptors (on the signaling cell) are integrated transmembrane proteins. Thus direct contact between two cells is required for a signaling event to occur. Upon binding to its ligand, the Notch receptor undergoes conformational changes, which expose a cleavage site for the enzyme *γ*-secretase. This enzyme releases the Notch intracellular domain (NICD) from its connection to the plasma membrane. The NICD is then transported to the nucleus, where it activates the CSL transcription factors and thus initiates the expression of several other downstream transcription factors, including Hes, Hey, Snail, as well as cyclins, and other genes (9, 10). Since the proteolytic cleavage of the Notch receptor plays a crucial role in the function of the signaling pathway, targeting the *γ*-secretase activity with small-molecule inhibitors, such as DAPT, has become a commonly used strategy to study the pathway.

Even though Notch signaling has been extensively studied in the context of embryonic development, cancer, and stem cell function, much less is known about the role(s) that this pathway plays in post-traumatic regeneration. The available data suggests that the Notch signaling is crucial for regeneration in several model organisms. For example, it has been shown that Notch is required for proper head and tentacle regeneration in *Hydra* (11), control of cell differentiation during retinal regeneration in newts (12), regulation of cardiomyocyte proliferation in zebrafish heart regeneration (13), and whole-body regeneration in planarians (14). Nevertheless, the function of the pathway is poorly understood in other regenerative species, such as echinoderms.

Echinoderms are a phylum of multicellular animals with highly regenerative species that can regrow almost all tissue types. Echinoderms share a deep common evolutionary ancestor with chordates. These features make echinoderms particularly attractive model organisms in regenerative biology (15, 16). However, the molecular mechanisms that drive echinoderm regeneration, including the functional role(s) of the key signaling pathways, are still poorly understood at the mechanistic level. Genes encoding all major components of the Notch pathway are present in echinoderms. They have been first identified in the fully sequenced genome of the sea urchin *Strongylocentrotus purpuratus* (Stimpson, 1857) (5). Similarly, our earlier data from the transcriptomic analysis showed that key members of the Notch signaling pathway were expressed in both the uninjured and regenerating radial nerve cord of the sea cucumber *Holothuria (Selenkothuria) glaberrima* Selenka, 1867 (17). These genes include the Notch receptor, ligands (Delta and Serrate), the transcriptional regulator RBPJ, two Notch target genes of the Hes family, and the Notch signaling modulator Numb. The only context, in which expression of those genes was studied at the cell and tissue levels in echinoderms, was sea urchin embryogenesis (18, 19). The only functional study of the Notch signaling pathway in the context of adult echinoderm regeneration was performed in the sea urchin *Lytechinus variegatus* (20). This work demonstrated the requirement of the functional Notch signaling for the proper outgrowth of amputated external appendages, such as spines and podia of *Lytechinus variegatus* (Lamarck, 1816). The cellular and molecular processes regulated by Notch signaling in echinoderm regeneration remain unknown. In addition, echinoid spines and podia are relatively simple structures. The role of the Notch signaling pathway in the regeneration of more complex organ systems and appendages in adult echinoderms has yet to be addressed.

Our aim in this study is to establish the functional role of the Notch signaling in arm regeneration in the brittle star *Ophioderma brevispina* (Say, 1825) and identify the target genes that are regulated by the pathway. Brittle star arms are segmented body appendages with complex internal anatomy. Each brittle star arm contains a calcareous endoskeleton composed of serial vertebral ossicles and several peripheral elements. Associated with the skeleton, the brittle star arm has a system of muscles and ligaments, two systems of coelomic canals, and a complex nervous system including a radial nerve and numerous peripheral nerves (21, 22). Brittle stars have emerged as important models in regenerative biology. They have been used in studies of skeletogenesis and biomineralization (23, 24), morphogenesis, and regulation of growth and differentiation (25). Here, we show that exposing regenerating brittle stars to the Notch pathway antagonist DAPT significantly impairs regeneration. We also identified target genes regulated by the pathway by performing a transcriptome wide gene expression analysis. We show that Notch regulates a multitude of biological processes involved in arm regeneration, including the extracellular matrix composition and remodelling, cell proliferation, death and migration, activity of mobile genetic elements, and the innate immune response. Our data also indicate an extensive cross-talk between Notch signaling and other key cell signaling pathways, such as Wnt, TGF-*β*, Toll, JNK and others, indicating that regeneration depends on a complex regulatory environment.

## Results

### The Notch signaling pathway in the brittle star

As expected, genes corresponding to the major components of the Notch pathway (i.e., the Notch receptor, the Delta and Serrate ligands, the transcriptional regulator RBPJ, two Notch target genes of the Hes family, and the Notch signaling modulator), as well as a Notchless protein homolog that plays a role in the Notch signaling pathway, were identified in the complete transcriptome of *O. brevispina*. Details are provided on a general Feature Format version 3 (GFF3) file, available in the supplements to this article (File S1).

### Pharmacological inhibition of the Notch signaling slows down arm regeneration

On day 14 post-autotomy, after continuous exposure of the treatment and control cohorts to the DAPT reagent (3 *µ*M) or DMSO (vehicle), respectively, two arms from each individual were processed for histological analysis. Six individuals per experimental group were used. As compared to the control individuals, the length of the regenerated arm (outgrowth) in the DAPT-treated cohort was on average 37% shorter (967 *µ*m vs 605 *µ*m, T-test *p*-value = 0.002) (Fig. 1, 2A). Likewise, the newly regenerated arm portion in the DAPT-treated animals had, on average, two fewer segments then the arms in the control group (5.5 vs 7.5, T-test *p*-value = 0.047) (Fig. 2B). Even though the new arms in the DAPT-treated individuals were smaller and less segmented, they still contained all major radial organs, including the radial nerve cord, the radial canal of the water-vascular system, and the arm coelom (compare Fig. 1C and D)

**Fig. 1.**
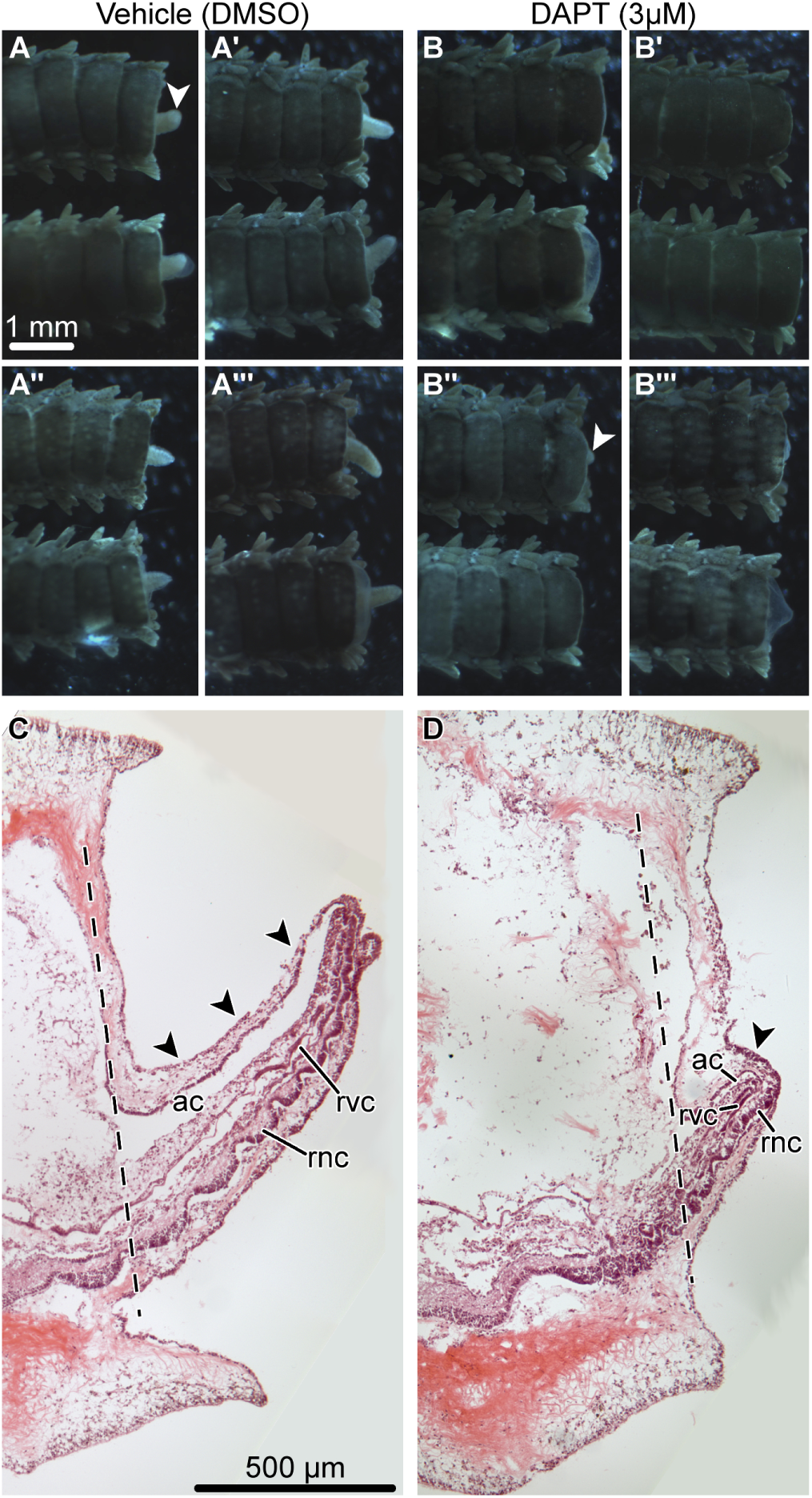
Effect of DAPT (3 *µ*M) treatment on the structure of the regenerating arm (day 14 post-autotomy). A – A‴: Aboral view of regenerating arms from four representative control animals exposed to DMSO (vehicle). Two arms from each animal are shown. B – B‴: Aboral view of regenerating arms from four representative animals exposed to 3 *µ*M DAPT. Two arms from each animal are shown. C and D: Representative sagittal sections through the regenerating arm of a control (DMSO-treated) individual (C) and a DAPT-treated individual (D). Hematoxylin and eosin staining. *Arrowheads* show the arm outgrowth (regenerate). *Dashed lines* show the position of the autotomy plane. Abbreviations: *ac* – arm coelom; *rnc* – radial nerve cord; *rvc* – radial canal of the water-vascular system.

**Fig. 2.**
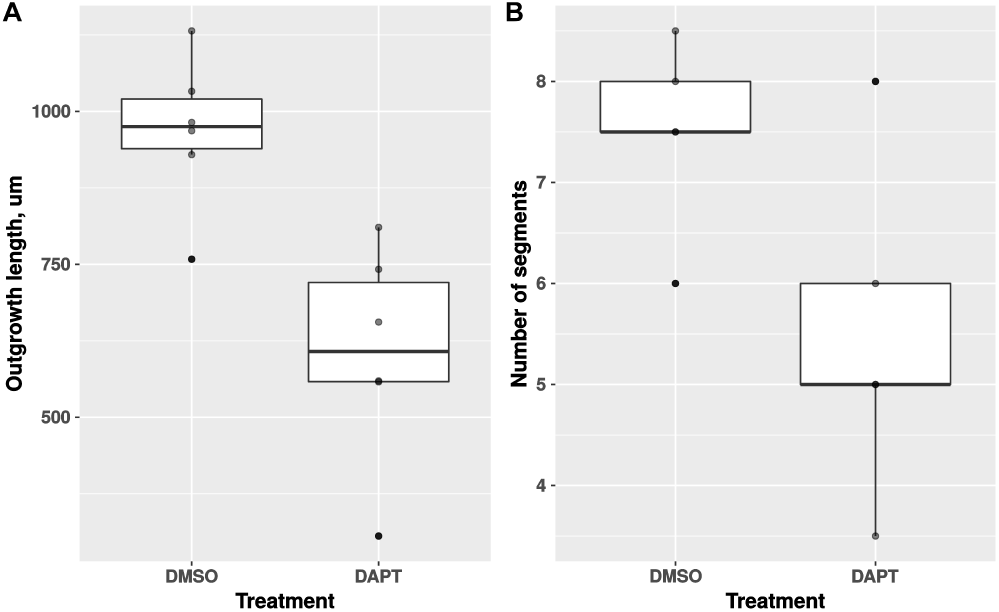
Boxplots showing the effect of DAPT (3 *µ*M) treatment on the length of the outgrowth (A) and the number of segments in the regenerating arm (B). By day 14 post-autotomy, the DAPT treatment significantly reduced both the total length of the outgrowth (by the factor of 1.6, T-test *p*-value = 0.002) and the number of segments in the new arm (by the factor of 1.4, T-test *p*-value = 0.047).

### *De novo* transcriptome assembly

We are currently working on generating ‘omic resources for the brittle star *O. brevispina*. However, since at the time of writing the assembly and annotation of the full genome for this species is still in preparation, we chose to use a *de novo* assembled transcriptome as a reference to characterize the Notch pathway target genes.

The *de novo* transcriptome was generated from 17,318,775 MiSeq and 832,245,006 HiSeq quality filtered and adapter trimmed reads. The single MiSeq library represented pooled samples from intact and regenerating arms at different states of regeneration, whereas six HiSeq libraries corresponded to three control (DMSO-treated) and three DAPT-treated regenerating individuals on day 14 post-autotomy (see *Methods*). Sequence reads were assembled with Trinity (26, 27) into 2,463,269 contigs (1,169,021 Trinity “genes”) with the average/median contig length of 421.6/260 nt and contig N50 of 527 nt. The key assembly metrics are listed in Table 1.

**Table 1.** Key metrics of the *de novo* assembly.

| Metric | Value |
| --- | --- |
| Total assembled bases | 1,038,536,368 nt |
| Number of contigs | 2,463,269 |
| Number of Trinity “genes” | 1,169,021 |
| Shortest contig | 151 nt |
| Longest contig | 48,146 nt |
| Average contig length | 421.6 nt |
| Median contig length | 260 nt |
| Contig N50 | 527 nt |
| Overall read alignment rate | 94.09% |
| Reads aligned as proper pairs | 75.42% |
| Full-length transcript representation | 7,397 (out of 35,786) |
| Complete (single-copy/duplicated) BUSCOs | 98.7% (28.8%/69.9%) |

To assess the quality of the assembled transcriptome, we performed a series of benchmark tests. First, to assess the representation of reads in the transcriptome, all cleaned Illumina reads were aligned back to the assembled contigs with Bowtie 2 (28). The vast majority of the reads (94.09%) mapped back to the assembly. Of these, 75.42% of the reads aligned as proper pairs one (21.62%) or more (53.80%) times. Second, to determine the representation of completely or almost completely (>80% of the length) assembled protein-coding transcripts, we compared our contigs to the reference proteome of the sea urchin *S. purpuratus* (29), the echinoderm species with the best-annotated genome to date. This analysis showed that 7,397 sea urchin orthologs (out of 35,786) are represented in our transcriptome by full-length and nearly full-length transcripts.

Third, the completeness of the assembly in terms of protein-coding gene content was assessed using BUSCO (30) and the conserved metazoan gene dataset. Out of 978 genes (or 98.7%) in the metazoan database, 966 genes were recovered in the assembled transcriptome as “complete” (i.e., their length fell within two standard deviations of the BUSCO group mean length). Of these complete genes, 282 matched a single contig, whereas multiple copies represented the remaining 684. The high number of “duplicated” genes is a known phenomenon in *de novo* transcriptome assembly, as even in the absence of any sequencing errors, inherent biological complexity of the transcriptome (e.g., single nucleotide polymorphism and alternative splicing) makes assembly algorithms report multiple isoforms for individual genes (31).

### Identification of the Notch pathway target genes by a transcriptome-wide gene expression analysis

Identification of direct and indirect targets of the Notch signaling pathway was performed by transcriptome-wide gene expression comparison between the DAPT-treated cohort and the control (vehicle-treated) animals. To reduce the redundancy of the *de novo* assembled transcriptome and to be able to perform transcript expression quantification at the “gene” level, we clustered the 2,463,269 Trinity contigs into 1,012,954 “clusters” using the Corset tool (32). Corset was applied after the reads representing the libraries from the control and DAPT-treated individuals were aligned back to the assembled transcriptome (33). Differential gene expression analysis was performed with the DESeq2 package (34), which identified 1,978 significantly up-regulated transcripts and 2,434 significantly down-regulated in the DAPT-treated cohort, as compared to control (DMSO-treated) individuals. For the purposes of this study, differentially expressed genes were defined as those whose expression in response to DAPT treatment changed by at least the factor of 1.5 in either direction and the associated *p*-value of the statistical test adjusted for multiple comparisons was less than 0.05 (Fig. 3).

**Fig. 3.**
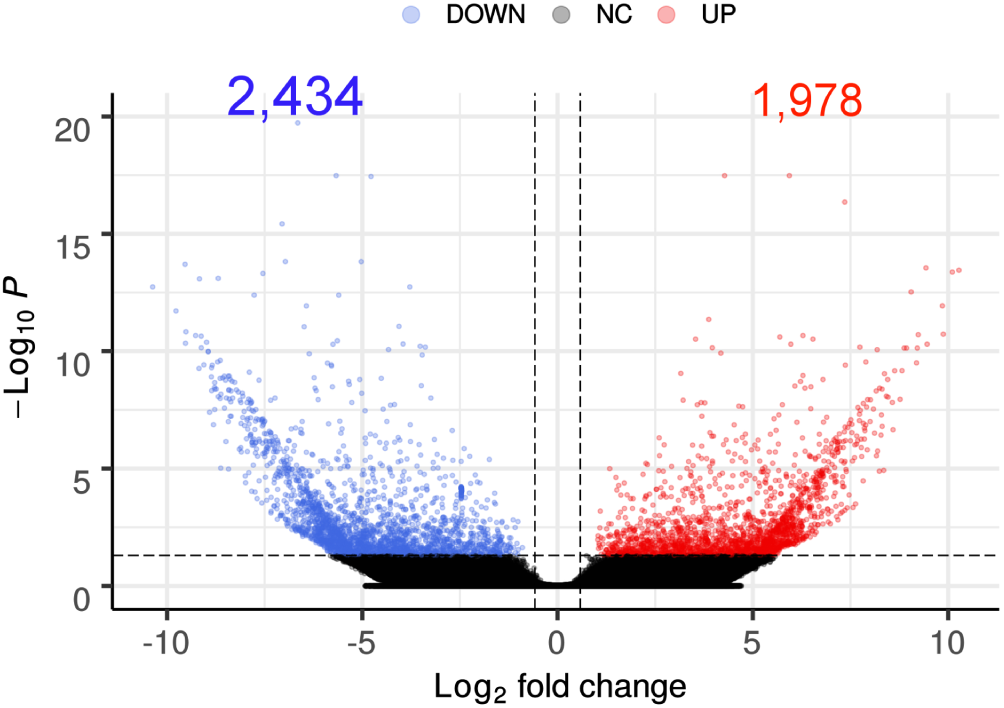
Differentially expressed genes in the DAPT-treated animals as compared to the control (DMSO-treated) cohort. Volcano plot showing the log2 fold change on the x-axis and -log10 of adjusted p-values on the y-axis. The significantly upregulated transcripts *(red)* are to the right, and significantly down-regulated transcripts *(blue)* are to the left. Each gene is represented with a dot. *Grey* dots represent the genes whose expression level did not change significantly in response to DAPT treatment. The differentially expressed contigs were defined as those whose associated adjusted *p-*value was less than 0.05 and the log2 fold change in expression exceeded ±0.58.).

To extract the biological meaning behind these extensive lists of genes, we used the DAVID functional annotation resource (35, 36). To this end, we annotated the transcripts representing the Corset clusters by parsing them against the Uniprot database. This approach yielded lists of Uniprot IDs for up- and down-regulated genes, containing 180 and 142 entries, respectively. As the “gene population background” (35), we used a list of 59,110 entries yielded by annotation of all Corset clusters. The percentage of the gene identifiers that were successfully “mapped” to the internal IDs within the DAVID knowledge base were 89%, 88%, and 92% for the list of down-regulated, up-regulated and “background” genes, respectively. In DAVID, we then used the “mapped” genes to perform functional annotation clustering by classifying the input genes into groups based on measuring relationships among annotation terms associated with these genes. Each cluster is assigned an enrichment score, which is defined as the geometric mean of the enrichment *p*-values associated with each annotation term in the group (35). The results of the DAVID cluster analysis are shown in Tables 2 and 3. The few genes, whose identifiers failed to map to the DAVID database, were annotated manually (Tables 4, 5).

**Table 2.**
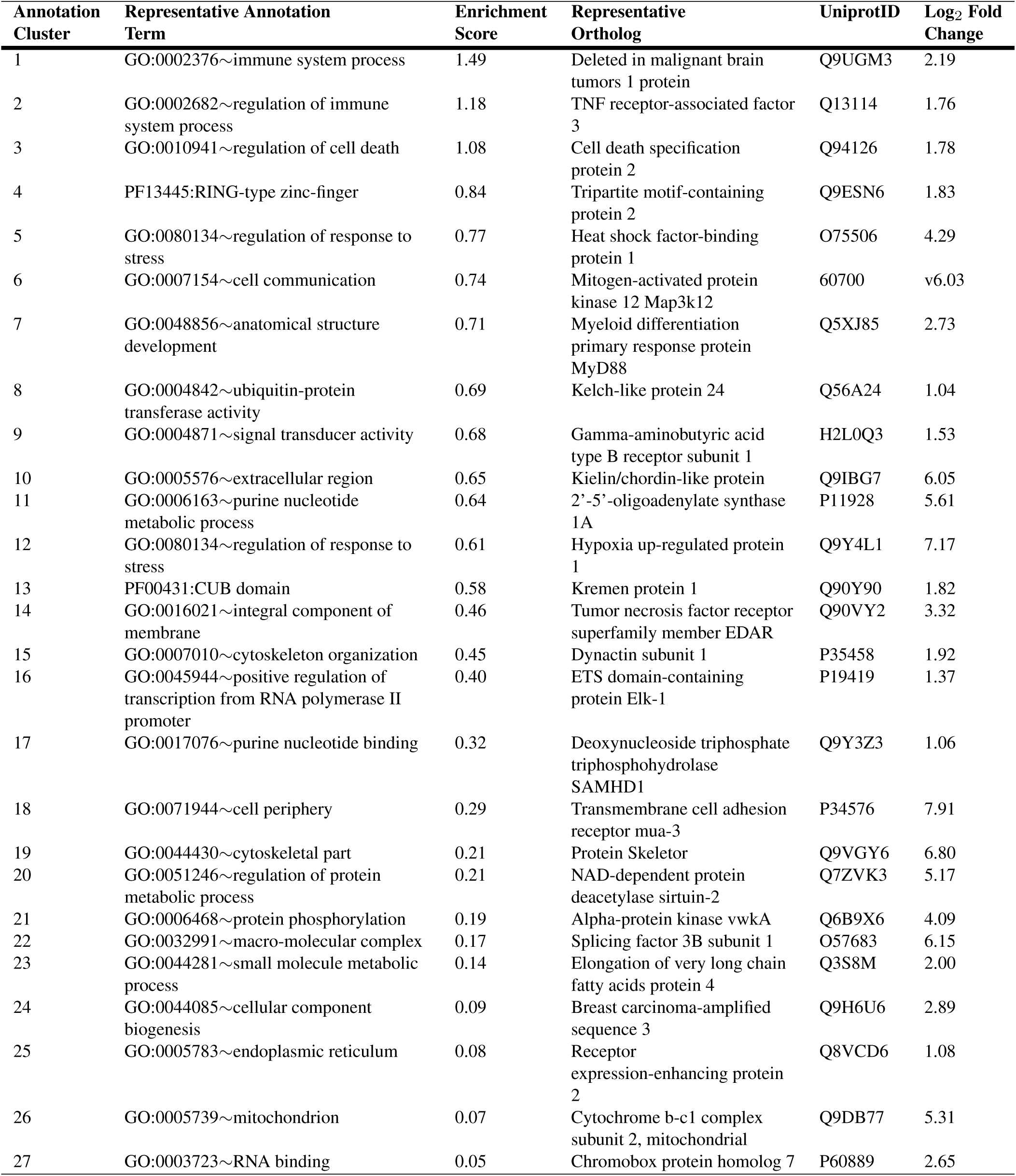
DAVID functional clustering of down-regulated genes.

**Table 3.**
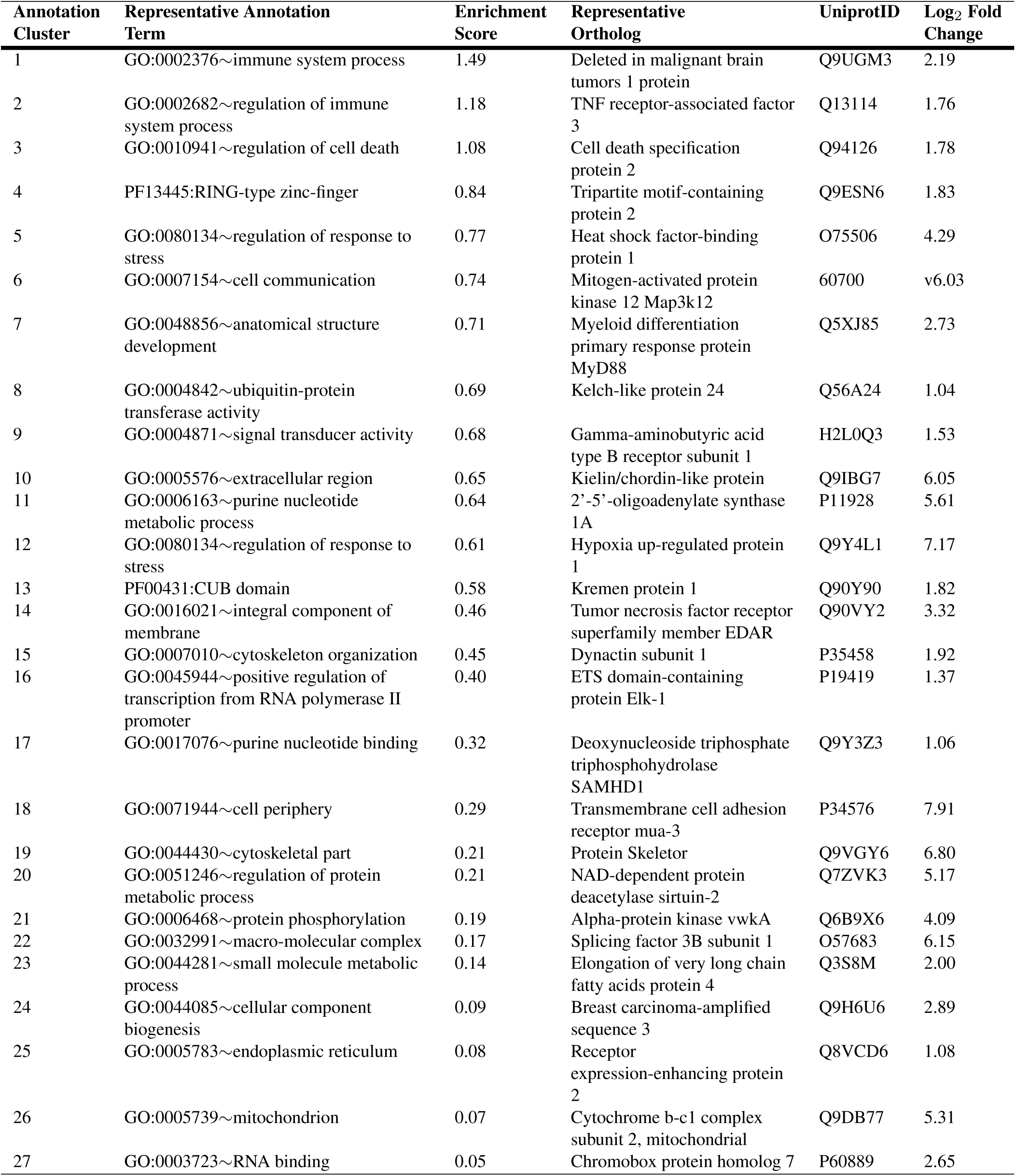
DAVID functional clustering of up-regulated genes.

**Table 4.**
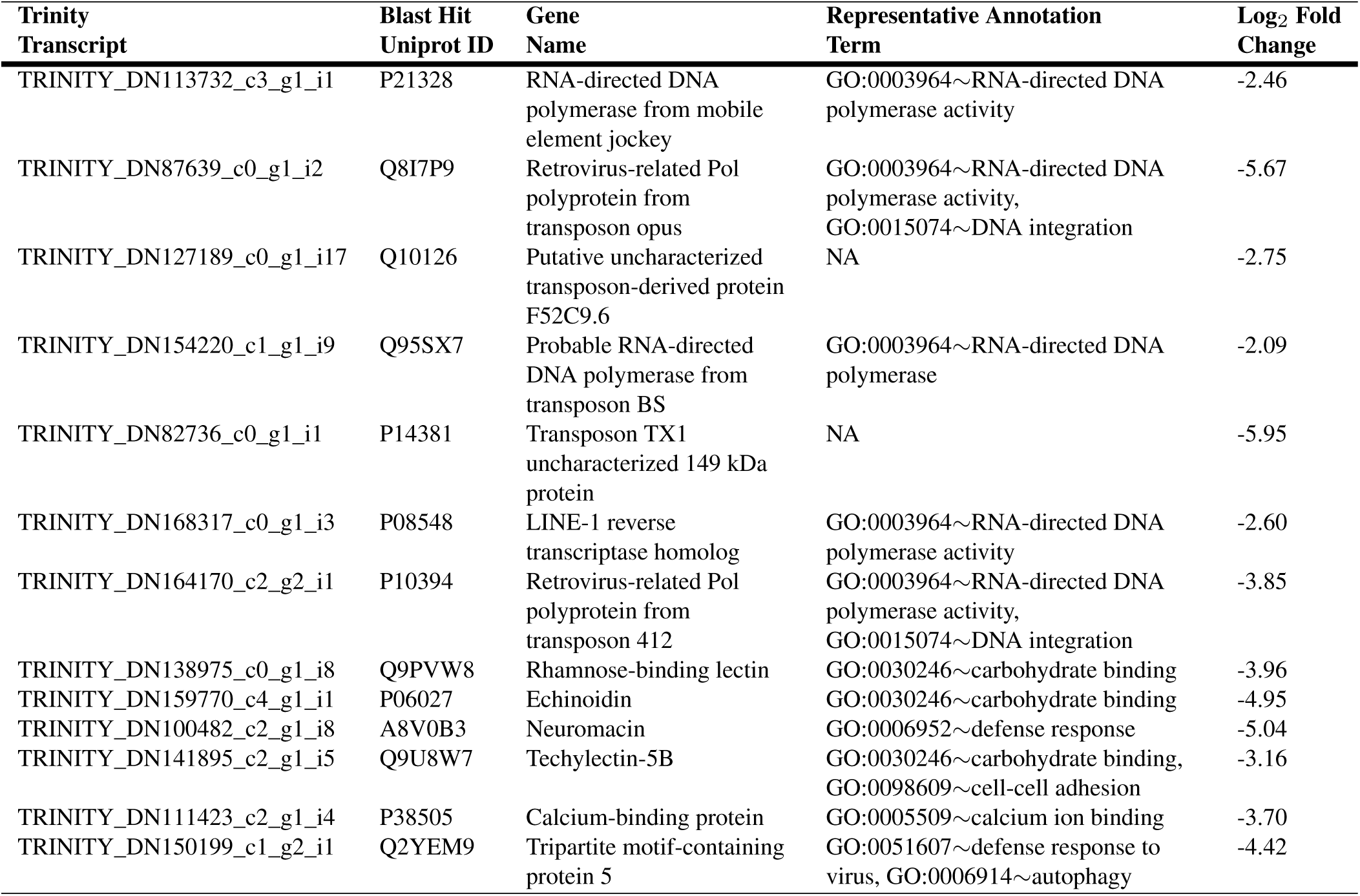
Down-regulated genes, unmapped by DAVID.

| Trinity Transcript | Blast Hit Uniprot ID | Gene Name | Representative Annotation Term | Log<sub>2</sub> Fold Change |
| --- | --- | --- | --- | --- |
| TRINITY_DN113732_c3_g1_i1 | P21328 | RNA-directed DNA polymerase from mobile element jockey | GO:0003964~RNA-directed DNA polymerase activity | -2.46 |
| TRINITY_DN87639_c0_g1_i2 | Q8I7P9 | Retrovirus-related Pol polyprotein from transposon opus | GO:0003964~RNA-directed DNA polymerase activity, GO:0015074~DNA integration | -5.67 |
| TRINITY_DN127189_c0_g1_i17 | Q10126 | Putative uncharacterized transposon-derived protein F52C9.6 | NA | -2.75 |
| TRINITY_DN154220_c1_g1_i9 | Q95SX7 | Probable RNA-directed DNA polymerase from transposon BS | GO:0003964~RNA-directed DNA polymerase | -2.09 |
| TRINITY_DN82736_c0_g1_i1 | P14381 | Transposon TX1 uncharacterized 149 kDa protein | NA | -5.95 |
| TRINITY_DN168317_c0_g1_i3 | P08548 | LINE-1 reverse transcriptase homolog | GO:0003964~RNA-directed DNA polymerase activity | -2.60 |
| TRINITY_DN164170_c2_g2_i1 | P10394 | Retrovirus-related Pol polyprotein from transposon 412 | GO:0003964~RNA-directed DNA polymerase activity, GO:0015074~DNA integration | -3.85 |
| TRINITY_DN138975_c0_g1_i8 | Q9PVW8 | Rhamnose-binding lectin | GO:0030246~carbohydrate binding | -3.96 |
| TRINITY_DN159770_c4_g1_i1 | P06027 | Echinoidin | GO:0030246~carbohydrate binding | -4.95 |
| TRINITY_DN100482_c2_g1_i8 | A8V0B3 | Neuromacin | GO:0006952~defense response | -5.04 |
| TRINITY_DN141895_c2_g1_i5 | Q9U8W7 | Techylectin-5B | GO:0030246~carbohydrate binding, GO:0098609~cell-cell adhesion | -3.16 |
| TRINITY_DN111423_c2_g1_i4 | P38505 | Calcium-binding protein | GO:0005509~calcium ion binding | -3.70 |
| TRINITY_DN150199_c1_g2_i1 | Q2YEM9 | Tripartite motif-containing protein 5 | GO:0051607~defense response to virus, GO:0006914~autophagy | -4.42 |

**Table 5.**
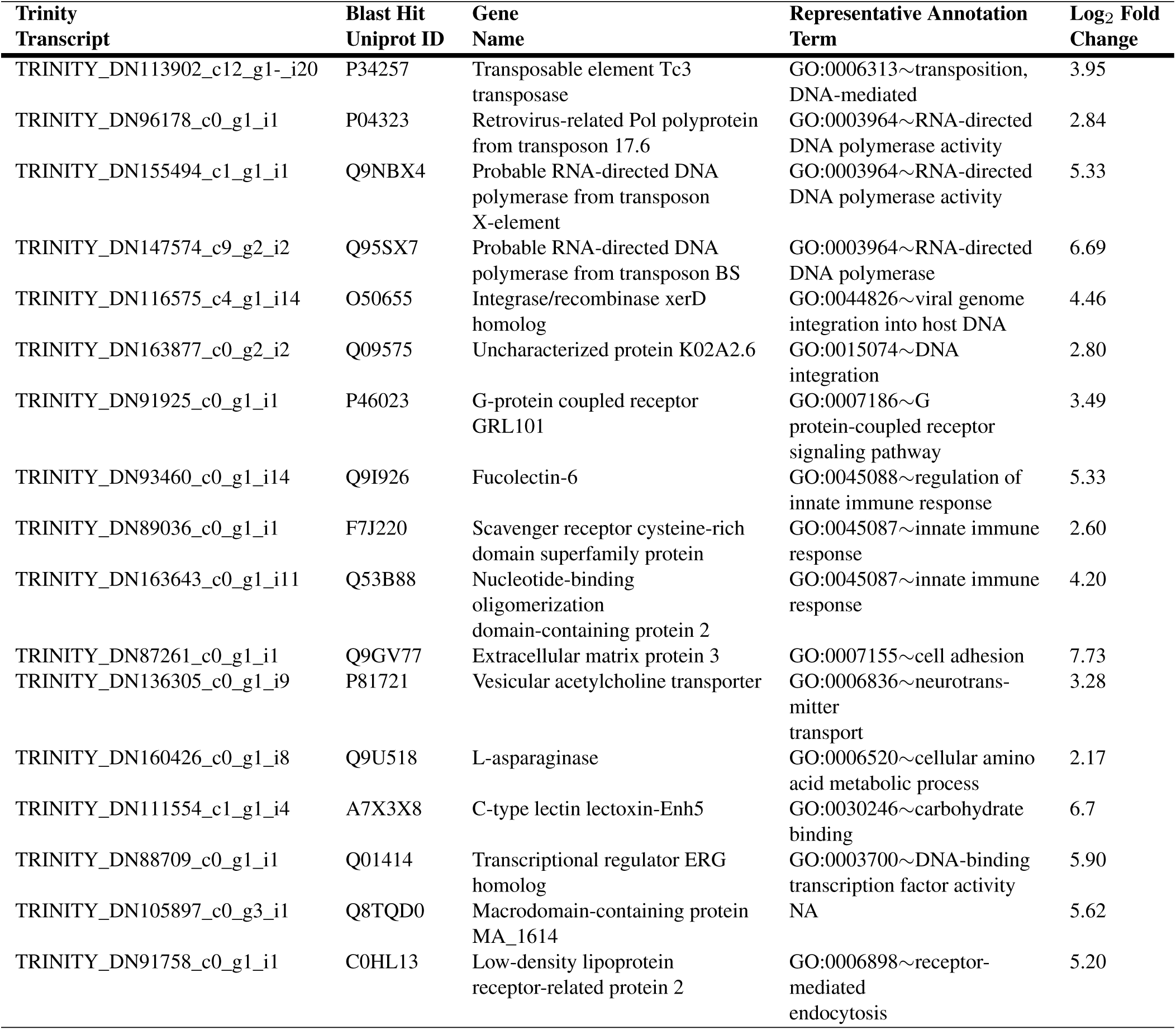
Up-regulated genes, unmapped by DAVID.

The down-regulated genes clustered by DAVID include genes that (a) code for extracellular matrix proteins and mediate mediate interactions between cells and the extracellular matrix (e.g., cell adhesion), (b) regulate multiple signaling pathways (including TGF-*β*, retinoic acid, JNK, p38, NF-kappa-B, JUN, TNF, Wnt, Toll-like receptor signaling, and EFGR signaling), (c) regulate innate immune response (antiviral response, phagocytosis, lectin pathway), (d) regulate protein ubiquitination involved in protein homeostasis, quality control, immunity, cell death, and regulation of cell signaling transduction, (e) cell survival, proliferation, and differentiation; (f) regulate semaphorin signaling, cytoskeleton organization/remodeling and cell migration, (g) regulate function of the endomembrane system, including the endoplasmic reticulum; (h) regulate cellular homeostasis, including regulation of intracellular ion concentration and ribosomal structure and function (Table 2).

Out of 14 down-regulated genes that did not get mapped to the DAVID database, seven represented proteins derived from retrotransposon genes (e.g., DNA-polymerase, reverse transcriptase, Pol). The remaining genes encode for three lectins, an antibacterial protein neuromacin, and the antiviral tripartite motif-containing protein 5 (Tables 4).

The up-regulated gene in DAVID functional clusters are known to be involved in (a) innate immune activity (antiviral and antibacterial), (b) cell signaling pathways (Toll-like, NF-kappa-B, JNK, p38, MAPK, VEGF, TOR, TGF-beta, BMP, Wnt), (c) positive regulation of neuronal and synaptic differentiation, (d) cell proliferation and differentiation, (e) cell death and removal of apoptotic cells by phagocytosis, (f) stress response (e.g., hypoxia, mitochondrial stress, negative regulation of the heat shock response), (g) epigenetic regulation (e.g., maintaining the transcriptionally repressing state by Polycomb proteins), (f) cell adhesion and migration, (g) RNA maturation, processing and splicing, (h) mitochondrial function in cellular respiration and cell death (Table 3). One of the genes that are up-regulated in response to inhibition of the Notch pathway is E3 ubiquitin-protein ligase Neurl1, an inhibitor of the Notch pathway, suggesting a control mechanism through mutual inhibition.

Among 18 up-regulated genes that did not have a match in the DAVID database, seven represented transcripts derived from transposable elements, including a transposase from a Tc3-like DNA transposon, reverse transcriptase genes from three retroelements, and two putative retrotransposon integrase genes. The remaining unmapped genes are involved in innate immune response, cell signaling, cell adhesion, lipoprotein endocytosis, and mino acid metabolism. In addition, one of the transcripts matched an ERG-like transcription factor (known protooncogene) (Table 5).

## Discussion

Here we show that the proper function of the Notch signaling pathway is required for arm regeneration in a brittle star. Pharmacological inhibition of the pathway with DAPT antagonist during the first 14 days of regeneration resulted in a significantly reduced overall length of the outgrowth and a smaller number of segments in the new arm. A similar effect of Notch pathway inhibition on regeneration was previously shown for sea urchin spines and podia. Treatment of DAPT resulted in a dosage-dependent reduction of regrowth of those structures post-amputation (20). This suggests that the involvement of Notch signaling in the regeneration of body appendages is conserved among echinoderm classes.

The transcriptome-wide quantitative comparison of gene expression between the control individuals and the individuals with the inhibited Notch pathway provided us with an insight into the biological roles of the pathway in the brittle star arm regeneration. This analysis shows that the Notch signaling pathway directly or indirectly affects more than two thousand downstream genes.

Functional annotation of the genes that significantly changed their expression in response to the pathway perturbation suggested that Notch controls a wide array of biological phenomena in regeneration. Additionally, this work provides the first evidence suggesting that the activity of some of the mobile DNA elements (transposons) is regulated by Notch signaling.

First, the Notch pathway regulates the expression of multiple components of crucial other cell signaling pathways in both positive and negative ways. This data indicates that individual pathways are not working in isolation during brittle star arm regeneration, but rather involved in a complex cross-coordination. This conclusion is consistent with previous studies that established the interaction of Notch signaling with other signaling pathway including, for example, the Bmp and Wnt pathways in the vertebrate heart development and repair (10), interaction between the Notch and Wnt signaling systems in *Drosophila* development and human cancers (37), and with the Wnt, BMP, and Yap/Taz pathways in regulation of neuronal stem cells (38).

Second, a number of studies have previously demonstrated a crucial role of the extracellular matrix (ECM) remodeling in echinoderm regeneration (24, 39–41). However, the upstream molecular mechanisms that initiate and coordinate these changes have remained unknown. Here, we show that in brittle star arm regeneration, Notch signaling is involved in both positive and negative regulation of different aspects of ECM composition and cell-ECM interaction, including cell migration.

Third, Notch positively regulates the innate immune pathways in regenerating tissues. These include lectins, antiviral responses, and phagocytosis. This elevated immune response might represent a reaction to the invasion of pathogens and the accumulation of damaged or necrotic cells at the site of autonomy. Alternatively, molecular components of the innate immune response are also known to function as key modulators of regeneration, including wound healing and cell division (42). For example, innate immune mechanisms are known to facilitate changes in epigenetic states and activate mechanisms required for nuclear reprogramming and cell dedifferentiation (43).

Fourth, the Notch signaling performs its classical functions in regulating cell proliferation, survival, programmed cell death, and differentiation in brittle star arm regeneration.

Fifth, the Notch pathway controls some of the cellular housekeeping functions, such as regulation of the endoplasmic reticulum, ion balance, ribosomes, RNA processing, and mitochondrial function.

Sixth, Notch regulates the transcriptional activity (both positively and negatively) of mobile genetic elements (RNA and DNA transposons). Transposons, and especially RNA transposons, are known to be active not only in the germ line, but also in somatic cells. Transposons affect the host cell gene expression via various mechanisms, including (1) providing promoter and enhancer sites that change the expression of the host genes, (2) creating splice and polyadenilation sites, and (3) through RNA interference (44). Although, in some cases, transposon activation is known to cause genetic disorders and autoimmune reactions (44), however, they also play important “positive” roles in adult and developing tissues. For example, in mammalian neurogenesis retrotransposition activity of LINE-1 elements contributes to neuronal diversity (45, 46). We have also recently shown that retrotransposons are differentially expressed in the sea cucumber neural regeneration. In response to injury, some of the retroelements increased their levels of expression, but there were others that were suppressed. The cells with activated retroelement activity remained alive and contributed to the newly regenerated tissues (47, 48).

We have hypothesized that mobile DNA plays a role in posttraumatic regeneration in echinoderms. The host molecular mechanisms that differentially regulate the transcriptional activity of mobile genetic elements are largely unknown. This work provides the first evidence suggesting that the activity of mobile DNA is controlled by a cell signaling pathway. Our data also provide further mechanistic insight into how Notch can positively regulated expression of retroelements. One of the genes repressed by active Notch signaling is deoxynucleoside triphosphate triphosphohydrolase SAMHD1 (Table 3), an antiviral protein that is also known to inhibit endogenous retroelements such as LINE-1 (49). Therefore, in the brittle star arm regeneration, the Notch pathway can activate retrotransposons through the double negative regulation mechanism.

Our data also provide an insight into how the Notch signaling pathway, once activated, might sustain its function in the brittle star arm regeneration. One of the genes that we found to be suppressed by the pathway is a homolog of Neuralized1, which is an antagonist of the Notch pathway (50).

## Conclusions

- The activity of the Notch signaling pathway is required for proper post-autotomy regeneration of the brittle star arm. Inhibition of the pathway results in a significant reduction of the outgrowth rate.
- There is an extensive cross-talk between the Notch pathway and other signaling pathways in the cell.
- The Notch pathway controls a wide range of cellular and molecular processes in the regenerating tissues, including:
  – composition and remodeling of the extracellular matrix
  – cell adhesion and migration
  – innate immune response
  – cell proliferation, survival, differentiation, and programmed cell death
  – “housekeeping functions” (functions of the endoplasmic reticulum, ribosomes, mitochondria, RNA processing, ion balance).
- The Notch pathway regulates (positively or negatively) the activity of several RNA and DNA transposable elements.
- A possible mechanism by which Notch sustains its activity in the regenerating tissues is suppression of Neuralized1.

## Methods

### Animal maintenance and pharmacological treatment

Adult individuals of the brittle star *Ophioderma brevispina* (Say, 1825) were purchased from the Marine Biological Laboratory (Woods Hole, MA) and were allowed to acclimate overnight in aerated sea water before being subjected to experimental procedures.

The DAPT reagent, N-[N-(3,5-Difluorophenacetyl)-L-alanyl]-S-phenylglycine t-butyl ester, was purchased from Sigma-Aldrich (catalog number D5942) and dissolved in DMSO (Dimethyl sulfoxide) to make a 20 mM stock solution.

The animals were divided into two cohorts (5 animals in each) – the DAPT treatment group and the control group. The DAPT treatment cohort was continuously exposed to 3 *µ*M DAPT prepared by diluting the stock solution in filtered seawater. The control group was exposed to the matching concentration of DMSO (vehicle). The animals were kept in glass vials submerged in 200 ml of the DAPT or DMSO solution, respectively. The solutions were changed daily.

The treatments started when the animals still had intact arms; then, after 24 hours, the arms were autotomized as described below, and the treatments continued until 14 days post-autotomy.

In each individual, all five arms were autotomized by squeezing a single arm segment with a fine forceps. Since the rate of regeneration in serpent stars is known to vary with the position of the autotomy plane along the proximodistal axis of the arm (25), we kept the injury paradigm consistent by autotomizing all arms at the level of the 15th segment (counting from the disk).

On day 14 post-autotomy, three of the five arms of each animal were pooled and used for total RNA extraction. The two remaining arms from each individual were processed for histological analysis.

### Histology and image analysis

Tissue samples were fixed overnight in 4% paraformaldehyde prepared in 0.01M PBS (pH 7.4) at 4°C. Fixed tissues were washed in PBS, decalcified in 10% EDTA, cryoprotected in graded sucrose solutions, and embedded in the Tissue-Tek OCT medium (Sakura). Serial longitudinal sections (10 *µ*m thick) were cut with a Leica CM1860 cryostat, collected on gelatin-covered slides, and incubated at 42°C overnight. They were then stained with Hematoxylin and Eosin and mounted in the DPX Medium (Electron Microscopy Sciences). The sections were photographed using an Olympus BX60 compound microscope equipped with a SPOT RT camera. The length of the regenerating arm was measured in six animals from each treatment group using the Fiji/ImageJ software (51, 52) in calibrated micrographs. The means between the two conditions were compared using Student’s T-test in R (53).

### Total RNA extraction

The tissue samples for RNA extraction included the most distal segment of the stump and the entire outgrowth (regenerate). The samples from three arms of each individual were pooled together and quickly homogenized in 500 *µ*l of ice-cold TRI reagent (Sigma-Aldrich, 93289) using a disposable Micro Tissue Homogenizer (Kimble, K7496250030). The homogenized samples were briefly vortexed, an additional 500 *µ*l of the TRI reagent were added to each tube, and the subsequent steps of RNA purification were performed as directed by the manufacturer’s protocol.

### Sequencing

#### Illumina HiSeq

Three total RNA samples from DAPT-treated individuals (each sample representing a different individual) and three samples from control individuals (DMSO-treated) were used for sequencing on the Illumina HiSeq 2000/2500 platform. The libraries were barcoded, pooled, and sequenced in a single flow cell in the Rapid Mode (2 × 100 bp). These reads were used both for the *de novo* transcriptome assembly and for gene expression analysis, as reads corresponding to each individual library could be identified and analyzed separately.

#### Illumina MiSeq

To facilitate the *de novo* assembly, we also prepared the following single combined sample to maximize the representation of expressed genes in the assembled reference transcriptome. Total RNA was extracted as above from intact animals and from untreated regenerating individuals on days 1, 3, 5, 18, and 30 post-autotomy. Each condition was represented by seven individuals. These 42 RNA samples (6 conditions × 7 individuals) were mixed in equimolar quantities and sequenced on the Illumina MiSeq platform (2 × 250 bp).

The final stages of library preparation and sequencing were outsourced to the Duke Center for Genomic and Computational Biology. The raw sequencing reads were deposited at the NCBI Sequence Read Archive (SRA) under the accession number (**to be added before publication**).

### *De novo* transcriptome assembly and quality assessment

The code used for the transcriptome assembly and subsequent RNAseq data analysis can be found in the accompanying additional file (File S2). The raw Illumina reads were trimmed with Trim Galore (54) to remove adapter sequences and low-quality bases at the 3’-end (with the base quality < 20). If a cleaned read was shorter than 20 nt, the entire read pair was discarded. This procedure yielded 17,318,775 MiSeq read pairs (combined library with pooled samples from different arm regeneration stages) with the read length ranging between 20–250 nt. Individual HiSeq libraries representing control and DAPT-treated individuals contained 34,676,875 ± 5,960,462 (mean ± standard deviation) read pairs with sequence length ranging between 20–100 nt. The total number of cleaned HiSeq read pairs was 832,245,006. All cleaned Illumina reads, from both the MiSeq and HiSeq technologies, were pooled and assembled with Trinity (26, 27) with the minimum reported contig length of 150 nt. The assembled transcriptome is available at (**to be added before publication**). The quality of the assembly was assessed by running several tests. First, to assess the representation of reads in the *de novo* transcriptome, we mapped all quality trimmed Illumina reads back to the assembled contigs using Bowtie 2 (Version 2.1.0) (28). Second, we determined the number of completely or almost completely (>80% of the length) assembled protein-coding transcripts by comparing our contigs to the reference proteome of the sea urchin *S. purpuratus* (29). Third, the completeness of protein-coding gene representation in the transcriptome was assessed with BUSCO (v3.0.2) (30) run in the “transcriptome mode” against the evolutionary conserved metazoan gene set (metazoa_odb9, creation date: 2016-02-13, number of species: 65, number of BUSCOs: 978).

### Identification of the main components of the Notch signaling pathway

To identify the main components of the Notch signaling pathway in the transcriptome of *O. brevispina*, we performed BLAST analysis (tblastn, E-value cutoff of 1*e* − 5; (55)) using Notch-related genes from the published transcriptomes of *H. glaberrima* (17) and *S. purpuratus* (5), keeping all hits. Each transcript identified this way was considered a putative Notch-related gene. We then used the NCBI’s conserved domain database (CDD; (56)) to search for conserved domains that categorize the target genes, including the Notch receptor, ligands (Delta and Serrate), the transcriptional regulator RBPJ, two Notch target genes of the Hes family (orthologous to HES-1 and HES-4), and the Notch signaling modulator Numb.

### Differential gene expression analysis

The cleaned reads from each of the six Illumina HiSeq libraries representing the control and DAPT-treated animals were aligned back to the assembled contigs using the Salmon tool (version v0.13.1) (33) with the *–dumpEq, –validateMappings*, and *–hardFilter* options. We then used Corset (version 1.08) (32) to mitigate the redundancy issue inherent to *de novo* transcriptome assemblies. Corset clusters isoforms into “genes” based on the proportion of shared reads and expression patterns and thus outputs gene-level counts.

Quantitative gene expression analysis was performed using the DESeq R package (34). To optimize memory usage, the raw count matrix was pre-filtered by only keeping rows that contained 10 reads or more. After normalizing the data and fitting the model, the independent hypothesis weighting approach for *p*-value adjustment implemented in the IHW R package (57) was applied to optimize the statistical power of the analysis. To improve fold change estimates for the gene expression data, a DESeq2 log2 fold change shrinking algorithm was applied. It uses information from all genes to improve the estimates for genes expressed at low levels with high dispersion values but does not change the total number of genes that are identified as differentially expressed (34). Genes were considered differentially expressed if their adjusted *p*-value was below 0.05 and the fold change in expression level exceeded 1.5 in either direction.

Functional annotation of differentially expressed genes was performed using DAVID (Database for Annotation, Visualization, and Integrated Discovery) (35), which was accessed from within R interface using the RDAVIDWebService package (36). To prepare data for DAVID, we annotated the Trinity transcripts representing all Corset clusters by matching them to the Uniprot database using BLASTX with the cutoff e-value threshold of 10^−5^. The resulting list of Uniprot accession IDs was used as the background gene set for DAVID. The up-regulated and down-regulated genes were annotated in the similar manner before being imported into DAVID. For functional annotation, we set the following annotation categories: “GOTERM_BP_ALL”, “GOTERM_MF_ALL”, “GOTERM_CC_ALL”, “KEGG_PATHWAY”, “BIO-CARTA”, and “PFAM”. DAVID cluster reports were parsed and analyzed using a custom R function (File S2).

## Supplementary Material

**File S1 General Feature Format File.** This file contains a GFF3 file which includes the sequence data and annotation of the main componetns of the Notch signaling pathway in *O. brevispina*.

**File S2 R Markdown Document.** This file contains all shell and R code used for transcriptome assembly and statistical analysis. The file can be opened in RStudio or any text editor.

