## Supplementary material for "Active Notch Signaling is Required for Arm Regeneration in a Brittle Star": File S2: S2_File.Rmd

````
---
output: html_notebook
---
# Analysis of morphological data
## Outgrowth Length
The measurement data is stored in the file *OutgrowthLength.xls*. Importing the data into R:

```{r}
library(readxl)
Outgrowth_Length <- read_excel("OutgrowthLength.xls", col_names = TRUE)
```

Running Student't T-test:

```{r, collapse=TRUE}
t.test(Outgrowth_Length$DMSO, Outgrowth_Length$DAPT, paired = FALSE, var.equal = TRUE)
```

Finding the  fold difference in outgrowth length between the DMSO-treated and DAPT-treated groups:
```{r, collapse=TRUE}
mean(Outgrowth_Length$DAPT)/mean(Outgrowth_Length$DMSO)
```

Plotting the data:
```{r, fig.width=5, warning=FALSE, message=FALSE, error=FALSE}
library(tidyverse)
ggplot(stack(Outgrowth_Length), aes(x = ind, y = values)) +
  geom_boxplot(notch = FALSE) +
  geom_point(alpha = 0.5) +
  scale_x_discrete(name = "Treatment") +
  scale_y_continuous(name = "Outgrowth length, um") +
  theme(axis.text = element_text(face = "bold", size = 11),
        axis.title = element_text(face = "bold", size = 13)
        )
```
***

## Number of segments
Creating a data frame with the data:
```{r}
DMSO_segments <- c(6, 7.5, 7.5, 8.5, 8)
DAPT_segments <- c(6, 5, 8, 3.5, 5)
Segments <- data.frame(DMSO_segments, DAPT_segments)
```
Running Student's T-test:
```{r, collapse=TRUE}
t.test(Segments$DMSO_segments, Segments$DAPT_segments, paired = FALSE, var.equal = TRUE)
```
Finding the fold difference in the number of segments between the DAPT-treated and control cohorts:
```{r, collapse=TRUE}
mean(Segments$DAPT_segments)/mean(Segments$DMSO_segments)
```

Plotting the data:
```{r, fig.width = 5}
ggplot(stack(Segments), aes(x = ind, y = values)) +
  geom_boxplot(notch = FALSE) +
  geom_point(alpha = 0.5) +
  scale_x_discrete(name = "Treatment") +
  scale_y_continuous(name = "Number of segments") +
  theme(axis.text = element_text(face = "bold", size = 11),
        axis.title = element_text(face = "bold", size = 13)
  )
```
***

# De novo transcriptome assembly
1. The raw Illumina reads were cleaned using [Trim Galore](https://www.bioinformatics.babraham.ac.uk/projects/trim_galore/) as follows:
```{bash, eval = FALSE, echo = TRUE, warning=FALSE, message=FALSE, error=FALSE}
trim_galore --paired --quality 20 --fastqc
```
This performs the following:

  + Trimming of low-quality bases at the 3'-end of each read (with the threshold Phred score of 20)
  + Removal of adapter sequences
  + Discarding sequences shorter than 20 bp. The `--paired` option removes the entire read pair if at least one of the two sequences is shorter than the threshold length

2. The cleaned reads were concatenated into two files (''left'' and ''right'') and assembled with [Trinity](https://github.com/trinityrnaseq/trinityrnaseq/wiki):
```{bash, eval = FALSE, echo = TRUE}
Trinity --seqType fq  --max_memory 890G \
--left /nobackup/echinotol/rnaseq/trinity/bigfile-brittle-R1.fq \
--right /nobackup/echinotol/rnaseq/trinity/bigfile-brittle-R2.fq \
--CPU 32 \
--min_contig_length 150	
```


# Assembly metrics and quality assessment
1. Running `TrinityStats.pl` on the `Trinity.fasta` file yields some basic numbers:
  
  + Total assembled bases: 1,038,536,368
  + Average contig length: 421.61 nt
  + Median contig length: 260 nt
  + Contig N50: 527 nt
  + Number of contigs: 2,463,269
  + Number of Trinity "genes": 1,169,021
  
2. Some additional info can be gained by runing the `seqstat` unitility (from the `biosquid` suite available through the Ubuntu repository) on the *Trinity.fasta* file:
```{bash, eval = FALSE, echo = TRUE}
seqstat -a Trinity.fasta > Trinity_seqstat.txt
```

  + Shortest contig: 151 nt
  + Longest contig: 48,146 nt

Importing the *Trinity_seqstat.txt* file into R, "cleaning" it and using the data to generate a contig length distribution histogram:
```{r, message=FALSE, error = FALSE, warning=FALSE}
library(readr)
TrinityContigLen <- read_delim(file = "Trinity_seqstat.txt", delim = " ", col_names = FALSE, skip = 7, col_types = "-cn------") 
TrinityContigLen <- TrinityContigLen[1:(nrow(TrinityContigLen) - 7), ]
colnames(TrinityContigLen) <- c("Contigs", "Length")
```

Plotting the data:
```{r}
ggplot(TrinityContigLen, aes(x = Length)) +
  geom_histogram(bins = 100) +
  scale_x_log10() +
  theme(axis.text = element_text(face = "bold", size = 11),
        axis.title = element_text(face = "bold", size = 13)
        ) +
  labs(x = "Length", y = "Counts")
```

3. To assess **read representation in the assembly**, we aligned all the quality-trimmed Illumina reads back to the assembled contigs unsing [Bowtie2](https://github.com/BenLangmead/bowtie2).
  + Building a `bowtie2` index:
```{bash, eval=FALSE, echo=TRUE}
module load bowtie2/2.2.9
input=Trinity.fasta
basename=Trinity
bowtie2-build $input $basename
```
  + Running the alignment:
```{bash, eval=FALSE, echo=TRUE}
module load bowtie2/2.2.9 samtools/1.9

reads_left=bigfile-brittle-R1.fq
reads_right=bigfile-brittle-R2.fq
basename=Trinity
outname=aligned.bam

bowtie2 -p 10 -q --no-unal -k 20 -x $basename -1 $reads_left -2 $reads_right \
2>align_stats.txt | samtools view -@10 -o $outname
```
The overall alignment rate was 94.09%. Of these, 75.42% reads aligned as proper pairs one (21.62%) or more (53.80%) times.

4. **Representation of full length protein-coding genes in the assembly.** To assess the number of completely or almost completely assembled transcripts representing protein-coding genes, we compared our contigs to the [NCBI collection](ftp://ftp.ncbi.nih.gov/genomes/Strongylocentrotus_purpuratus/protein/) of the sea urchin (*Strongylocentrotus purpuratus*) proteins.
  + Building a blastable database:
  ```{bash, eval=FALSE, echo=TRUE}
module load blast/2.5.0+
input=Sp_protein.fa
makeblastdb -in $input -dbtype prot
  ```
  + Running the blast search:
```{bash, eval=FALSE, echo=TRUE}
module load blast/2.5.0+
query=Trinity.fasta
database=Sp_protein.fa
output=Sp_blastx.outfmt6
evalue=1e-6

blastx -query $query \
-db $database \
-out $output \
-evalue $evalue \
-num_threads 16 \
-max_target_seqs 1 \
-outfmt 6
```
  + Running the `analyze_blastPlus_topHit_coverage.pl` utility that is included with Trinity:
```{bash, eval=FALSE, echo=TRUE}
module load trinity/2.4.0

query=Trinity.fasta
database=Sp_protein.fa
output=Sp_blastx.outfmt6
evalue=1e-6

/apps/pkg/trinity-2.4.0/util/analyze_blastPlus_topHit_coverage.pl $output $query $database
```
  Here is the content of the *Sp_blastx.outfmt6.hist* output file:

  hit_pct_cov_bin | count_in_bin | >bin_below
  ----------------|--------------|-----------
  100             | 5527         | 5527
  90	            | 1870	       | 7397
  80	            | 1436	       | 8833
  70	            | 1350	       | 10183
  60	            | 1453	       | 11636
  50	            | 1450	       | 13086
  40	            | 1572	       | 14658
  30	            | 1682	       | 16340
  20	            | 1794	       | 18134

The biological meaning of this analysis is that there are 7,397 sea urchin orthologs (out of 35,786) that are represented in our transcriptome by full-length and nearly full-length transcripts (>80% alignment coverage).

5. The **completeness of the assembled transcriptome** (in terms of the gene content) was also assessed with [BUSCO](https://busco.ezlab.org/). The comparison was run agins the "Metazoa" gene set (metazoa_odb9, creation date: 2016-02-13, number of species: 65, number of BUSCOs: 978)

```{bash, eval=FALSE, echo=TRUE}
module load augustus/3.3 blast/2.5.0+ hmmer/3.2.1 busco/3.0.2 

sequences=Trinity.fasta
database=metazoa_odb9
output=Ob_busco_metazoa

run_BUSCO.py \
-i ${sequences} \
-o ${output} \
-l ${database} \
-m tran \
--cpu 16
```

Below is the output of the BUSCO analysis:
C:98.7%[S:28.8%,D:69.9%],F:1.1%,M:0.2%,n:978

Category                        | Count
--------------------------------|------
Complete BUSCOs (C)             | 966
Complete single-copy BUSCOs (S) | 282
Complete duplicated BUSCOs (D)  | 684
Fragmented BUSCOs (F)           | 11
Missing BUSCOs (M)              | 1
Total BUSCO groups searched (n) | 978

***

# Gene Expression Analysis
## Read Mapping with [Salmon](https://github.com/COMBINE-lab/salmon)
* Building the Salmon index:

```{bash, eval=FALSE, echo=TRUE}
transcriptome=Trinity.fasta
salmon index --index Trinity --transcripts ${transcriptome} -k 31
```

* Quantifying the reads:

```{bash, eval=FALSE, echo=TRUE}
transcriptome=Trinity.fasta
reads=DirName

FILES=`ls ${reads}/*_R1.fq | sed "s|${reads}/||g" | sed 's/_R1.fq//g'` #Getting the names of the treatments
for F in ${FILES} ; do
reads1=${reads}/${F}_R1.fq
reads2=${reads}/${F}_R2.fq
salmon quant --index Trinity --libType IU --dumpEq --validateMappings --hardFilter -1 ${reads1} -2 ${reads2} --output ${F}.salmon.out
done
```

## Obtaining the gene-level counts by clustering the isoforms into "genes" with [Corset](https://doi.org/10.1186/s13059-014-0410-6):

```{bash, eval=FALSE, echo=TRUE}
corset -g DMSO,DMSO,DMSO,DAPT,DAPT,DAPT -n DMSO_1,DMSO_2,DMSO_3,DAPT_1,DAPT_2,DAPT_3 -i salmon_eq_classes DMSO_Animal?.salmon.out/aux_info/eq_classes.txt DAPT_Animal?.salmon.out/aux_info/eq_classes.txt
```
Corset outputs two files *clusters.txt* and *counts.txt*. The first file, contains cluster assignment data, whereas the second one lists the number of reads assigned to each cluster. This second file is then used as an input for differential gene expression analysis with DESeq2.

## Differential gene expression analysis with [DESeq2](https://doi.org/10.1186/s13059-014-0550-8):
### Preparing the data

* Creating the metadata dataframe (*samples*) with information about the samples:
```{r, eval=TRUE, echo=TRUE}
    condition <- rep(c("DMSO", "DAPT"), each = 3)
    type <- rep("paired-end", times = 6)
    samples <- data.frame(condition, type)
    rownames(samples) <- c(
      paste0(rep("DMSO_", 3), c(1:3)), 
      paste0(rep("DAPT_", 3), c(1:3))
    )
```

* Loading the raw count data:
```{r}
    count_data <- read.delim(file = "counts.txt", header = TRUE, row.names = 1)
```

* Making sure that the sample name and order in the metadata dataframe matches those in the raw count dataframe:
```{r, collapse=TRUE}
    all(rownames(samples) == colnames(count_data))
```

* Generating the DESeq2 data set:
```{r}
    library("DESeq2")
    dds <- DESeqDataSetFromMatrix(
      countData = count_data,
      colData = samples,
      design = ~ condition
    )
```

* Prefiltering -- keeping only the rows with at least 10 reads:
```{r}
keep <- rowSums(counts(dds)) >= 10
dds <- dds[keep, ]
```

* Normalizing the counts: estimating the size factor:
```{r}
  dds <- estimateSizeFactors(dds)
  sizeFactors(dds)
```

### Quality assessment of the count data:

* Applying variance stabilizing transformation (VST) to perform a log2 transformation of the normalized count data:

```{r, eval=TRUE, echo=TRUE}
vsd_data <- vst(dds, blind = TRUE)
```

* Extracting the matrix of transformed data:
```{r}
vsd_data_matrix <- assay(vsd_data)
```

* Computing the pairwise correlation values between the samples:
```{r}
vsd_data_correlations <- cor(vsd_data_matrix)
```

* Drawing a heatmap of sample-to-sample distances:
```{r, message=FALSE, warning=FALSE}
library(pheatmap)
library(dplyr)
pheatmap(vsd_data_correlations, annotation = select(samples, condition))
```

* Performing principal component analysis (PCA)
```{r}
plotPCA(vsd_data, intgroup = "condition")
```

### Differential expression analysis

* Fitting and testing the model:

```{r}
dds <- DESeq(dds)
```

* Plotting the dispersion estimates:

```{r}
plotDispEsts(dds)
```
DESeq2 makes the assumption that the dispersion (variance squared) generally decreases with increasing mean of normalized counts. It is generally the case in our data. The black dots are dispersion estimates for each individual gene. The red line is fitted to show how dispersion is dependent on the mean. Each gene's estimate is then shrunk towards the red line to yield the final estimates (blue dots) that are used in the hypothesis test.

* Extracting the results of the differential gene expression analysis. Applying [Independent Hypothesis Weighting approach](ttps://www.nature.com/articles/nmeth.3885) to optimize the statistical power of the analysis:
```{r}
library(IHW)
Res_IHW <- results(dds, contrast = c("condition", "DAPT", "DMSO"), filterFun = ihw)
```

MA plot for this dataset:
```{r}
plotMA(Res_IHW, alpha = 0.05, ylim = c(-15, 15))
```

* Shrinking the log2 fold changes (LFC).
To improve the fold change estimates for the data, DESeq2 allows to shrink LFC estimates. It uses information from all genes to improve the estimates for genes with low counts/high dispersion values. LFC shrinkage does not change the total number of genes that are identified as differentially expressed.

```{r}
Res_IHW_shrunken <- lfcShrink(dds, contrast = c("condition", "DAPT", "DMSO"), res = Res_IHW)
```

MA plot for this dataset:
```{r}
plotMA(Res_IHW_shrunken, alpha = 0.05, ylim = c(-10, 10))
```

```{r}
library(EnhancedVolcano)

colors <- rep("black", nrow(Res_IHW_shrunken))
names(colors) <- rep("NC", nrow(Res_IHW_shrunken))

colors[which(Res_IHW_shrunken$log2FoldChange >= 0.58 & Res_IHW_shrunken$padj < 0.05)] <- "red2"
names(colors) [which(Res_IHW_shrunken$log2FoldChange >= 0.58 & Res_IHW_shrunken$padj < 0.05)] <- "UP"

colors[which(Res_IHW_shrunken$log2FoldChange <= -0.58 & Res_IHW_shrunken$padj < 0.05)] <- "royalblue"
names(colors) [which(Res_IHW_shrunken$log2FoldChange <= 0.58 & Res_IHW_shrunken$padj < 0.05)] <- "DOWN"

EnhancedVolcano(
  Res_IHW_shrunken,
  lab = rownames(Res_IHW_shrunken),
  x = "log2FoldChange",
  y = "padj",
  pCutoff = 0.05,
  FCcutoff = 0.58,
  selectLab = "",
  colCustom = colors,
  colAlpha = 0.3,
  title = "",
  subtitle = "",
  caption = "",
  ylim = c(0, 20)
)
```

Extracting significant genes (with fold-change cut-off of 1.5 and padj cutoff of 0.05)
```{r}
library(dplyr)
library(tibble)
Res_Sig <- Res_IHW_shrunken %>%
  as.data.frame() %>%
  rownames_to_column(var = "gene") %>%
  filter(padj < 0.05 & abs(log2FoldChange) >= 0.58)
```

Exporting the data on differentially expressed genes into a csv file:
```{r, eval=FALSE, echo = TRUE}
write.csv(Res_Sig, file = "DE_genes.csv", quote = FALSE, row.names = FALSE)
```


Checking the number of up-regulated genes:
```{r}
nrow(
  filter(Res_Sig, log2FoldChange >= 0.58)
)
```

Checking the number of down-regulated genes:
```{r}
nrow(
  filter(Res_Sig, log2FoldChange <= 0.58)
)
```

### Annotation of differentially expressed genes:
Extracting the cluster IDs:
```{bash, eval = FALSE, echo = TRUE}
cat DE_genes.csv | cut -f1 -d',' | sed 1d > DE_cluster_IDs.txt
```

Extracting the sequences corresponding to the clusters using the [fetchClusterSeqs.py](https://github.com/Adamtaranto/Corset-tools) utility:
```{bash, eval = FALSE, echo = TRUE}
transcriptome=Trinity.fasta
ClustersOfInterest=DE_cluster_IDs.txt
MappingFile=clusters.txt
out=`echo ${ClustersOfInterest} | sed 's/_Clusters.txt//g'`

./fetchClusterSeqs.py -l -i ${transcriptome} -t ${ClustersOfInterest} -c ${MappingFile} -o ${out}.fasta
```

Running BlastX against the uniprot database:
```{bash, eval = FALSE, echo = TRUE}
query=DE_cluster_IDs.txt.fasta
database=uniprot_sprot.fasta
output=${query}.blastp.outfmt6
evalue=1e-5

blastx -query $query \
 -db $database \
 -out $output \
 -evalue $evalue \
 -num_threads 16 \
 -max_target_seqs 1 \
 -outfmt 6
```

Reading blast annotation into R:
```{r}
library(readr)
blast <- read_tsv(file = "DE_cluster_IDs.txt.fasta.blastp.outfmt6", col_names = FALSE)
head(blast)
```

Getting rid of unnecessary columns, leaving only query ID, blast hit ID, and the e-value:
```{r}
library(dplyr)
blast <- select(blast, 1, 2, 11)
head(blast)
```

Separating the first column into two: cluster names and transcript names:
```{r}
library(tidyr)
blast <- separate(blast, col = 1, sep = "_", into = c("cluster", "transcript"), extra = "merge")
head(blast)
```

Modifying the third column now, keeping the UniprotID of the blast hit only, renaming the last column:
```{r}
blast <- blast %>%
  separate(col = 3, sep = "\\|", into = c("x", "BlastHitUniprotID", "y")) %>%
  select(- c("x", "y"))
blast <- rename(blast, "BlastHitEvalue" = X11)
```

Resolving redundancy in the annotation dataset by keeping only one entry per cluster with the lowest e-value:
```{r}
blast <- blast %>%
  group_by(cluster) %>%
  filter(BlastHitEvalue == min(BlastHitEvalue))
```

Adding the blast annotations to the Res_Sig dataframe:
```{r}
Res_Sig_Annot <- left_join(Res_Sig, blast, by = c("gene" = "cluster"))
```

Extracting the annotated up- and down-regulated genes:
```{r}
Res_Sig_Annot_UP <- Res_Sig_Annot %>%
  drop_na() %>%
  filter(log2FoldChange >= 0.58)

Res_Sig_Annot_DOWN <- Res_Sig_Annot %>%
  drop_na() %>%
  filter(log2FoldChange <= 0.58)
```

###  Functional enrichment analysis with [DAVID](https://www.bioconductor.org/packages/release/bioc/html/RDAVIDWebService.html)

We need a background gene list for DAVID. Therefore, I am extracting all cluster IDs from the corset output:
```{bash, eval=FALSE, echo=TRUE}
clusters.txt | cut -f2 > cluster_ID_ALL.txt
```

Extracting the contig sequences representing the clusters:
```{bash, eval=FALSE, echo=TRUE}
transcriptome=Trinity.fasta
ClustersOfInterest=cluster_ID_ALL.txt
MappingFile=../clusters.txt
out=`echo ${ClustersOfInterest} | sed 's/\.txt//g'`

./fetchClusterSeqs.py -l -i ${transcriptome} -t ${ClustersOfInterest} -c ${MappingFile} -o ${out}.fasta
```

Annotating these sequences agains the Uniprot database:
```{bash, eval=FALSE, echo=TRUE}
query=cluster_ID_ALL.fasta
database=uniprot_sprot.fasta
output=${query}.blastp.outfmt6
evalue=1e-5

echo "Start:" && date

blastx -query $query \
 -db $database \
 -out $output \
 -evalue $evalue \
 -num_threads 16 \
 -max_target_seqs 1 \
 -outfmt 6
```

Extracting the gene lists for annotation:

* List of up-regulated genes:
```{r}
library(dplyr)
UP_ID_list <- pull(Res_Sig_Annot_UP, var = "BlastHitUniprotID")
```

* List of down-regulated genes:
```{r}
DOWN_ID_list <- pull(Res_Sig_Annot_DOWN, var = "BlastHitUniprotID")
```

* The background list:
```{r}
blast_ALL <- read_tsv(file = "cluster_ID_ALL.fasta.blastp.outfmt6", col_names = FALSE)
blast_ALL <- blast_ALL %>%
  select(1, 2, 11) %>%
  separate(col = 1, sep = "_", into = c("cluster", "transcript"), extra = "merge")
blast_ALL <- blast_ALL %>%
  separate(col = 3, sep = "\\|", into = c("x", "BlastHitUniprotID", "y")) %>%
  select(- c("x", "y")) %>%
  rename("BlastHitEvalue" = X11) %>%
  group_by(cluster) %>%
  filter(BlastHitEvalue == min(BlastHitEvalue))
BACKGROUND_ID_list <- pull(blast_ALL, var = "BlastHitUniprotID")
```

Loading the package RDAVIDWebService:
```{r}
library(RDAVIDWebService)
```

Creating a connection to the DAVID service. Note - if an error pops up, the solution can be found [here](https://stackoverflow.com/questions/31480579/r-david-webservice-sudden-transport-error-301-error-moved-permanently):
```{r}
david <-DAVIDWebService$new(email = "", url="https://david.ncifcrf.gov/webservice/services/DAVIDWebService.DAVIDWebServiceHttpSoap12Endpoint/")
setTimeOut(david, 100000)
```

Defining the foreground and background gene lists:
```{r}
UP_david <- addList(david, UP_ID_list, idType = "UNIPROT_ACCESSION", listName = "UP", listType = "Gene")
DOWN_david <- addList(david, DOWN_ID_list, idType = "UNIPROT_ACCESSION", listName = "DOWN", listType = "Gene")

BACKGROUND_david <- addList(david, unique(BACKGROUND_ID_list), idType = "UNIPROT_ACCESSION", listName = "all", listType = "Background")
```

Setting the current background list:
```{r}
setCurrentBackgroundPosition(david, 39)
```

Specifying the annotation categories:
```{r}
setAnnotationCategories(david, c("GOTERM_BP_ALL", "GOTERM_MF_ALL", "GOTERM_CC_ALL", "KEGG_PATHWAY", "BIOCARTA", "PFAM"))
```

Performing the analysis for the DOWN-regulated gene set:
```{r}
DOWN_FuncAnnotChart_david <- getFunctionalAnnotationChart(david)
getFunctionalAnnotationChartFile(david, "DOWN_FuncAnnotChart_david.txt")

DOWN_Clusters_david <- getClusterReport(david, type = "Term")
getClusterReportFile(david, fileName = "DOWN_Clusters_david", type = "Term")

DOWN_AnnotTable <- getFunctionalAnnotationTable(david)
getFunctionalAnnotationTableFile(david, fileName = "DOWN_AnnotTable.txt")

DOWN_GeneCategoriesRepot_david <- getGeneCategoriesReport(david)
```

Getting the summary information about the DAVID clusters:
```{r}
summary(DOWN_Clusters_david)
```

Performing the analysis for the UP-regulated genes:

```{r}
setCurrentGeneListPosition(david, 1)
UP_FuncAnnotChart_david <- getFunctionalAnnotationChart(david)
getFunctionalAnnotationChartFile(david, "UP_FuncAnnotChart_david.txt")

UP_Clusters_david <- getClusterReport(david, type = "Term")
getClusterReportFile(david, fileName = "UP_Clusters_david", type = "Term")

UP_AnnotTable <- getFunctionalAnnotationTable(david)
getFunctionalAnnotationTableFile(david, fileName = "UP_AnnotTable.txt")

UP_GeneCategoriesRepot_david <- getGeneCategoriesReport(david)
```

Define the function to retrieve the UniprotID from the DAID clusters of interest and get the corresponding gene expression values:
```{r}
RetrieveExprValuesForDavid <- function(ClusterReport = c("DOWN_Clusters_david", "UP_Clusters_david"), ClusterNo, TermRowNo, Annot = c("Res_Sig_Annot_DOWN", "Res_Sig_Annot_UP")) {
  query <- members(ClusterReport)[[ClusterNo]][TermRowNo, 6]
  query <- strsplit(str_replace_all(query, pattern = " ", repl = ""), split = ",")
  query <- unlist(query)
  out <- Annot[Annot$BlastHitUniprotID %in% query, ]
  return(out)
}
# WOKRING EXAMPLE:
# Extract cluster 1 of down-regulated genes:
# cluster(DOWN_Clusters_david)[[1]]
#
# Extract the gene information for the second annotation term in this cluster:
# RetrieveExprValuesForDavid(DOWN_Clusters_david, 1, 2, Res_Sig_Annot_DOWN)
```

Exploring the genes, whose ID DAVID was not able to map:
```{r}
DOWN_unmapped <- unique(DOWN_david$unmappedIds)
Res_Sig_Annot_DOWN[Res_Sig_Annot_DOWN$BlastHitUniprotID %in% DOWN_unmapped, ]

UP_unmapped <- unique(UP_david$unmappedIds)
Res_Sig_Annot_UP[Res_Sig_Annot_UP$BlastHitUniprotID %in% UP_unmapped, ]
```
````
